# Targeting the nociceptive somatosensory system with AAV9 and AAV2retro viral vectors

**DOI:** 10.1101/2021.05.25.445559

**Authors:** Alexander G.J. Skorput, Reshma Gore, Rachel Schorn, Maureen S. Riedl, Ezequiel Marron Fernandez de Velasco, Bailey Hadlich, Kelley F. Kitto, Carolyn A. Fairbanks, Lucy Vulchanova

**Affiliations:** Department of Neuroscience, University of Minnesota, Minneapolis, MN; Department of Pharmacology, University of Minnesota, Minneapolis, MN; Department of Pharmaceutics, University of Minnesota, Minneapolis, MN

## Abstract

Adeno-associated viral (AAV) vectors allow for site-specific and time-dependent genetic manipulation of neurons. However, for successful implementation of AAV vectors, major consideration must be given to the selection of viral serotype and route of delivery for efficient gene transfer into the cell type being investigated. Here we compare the transduction pattern of neurons in the somatosensory system following injection of AAV9 or AAV2retro in the parabrachial complex of the midbrain, the spinal cord dorsal horn, the intrathecal space, and the colon. Transduction was evaluated based on Cre-dependent expression of tdTomato in transgenic reporter mice, following delivery of AAV9 or AAV2retro carrying identical constructs that drive the expression of Cre/GFP. The pattern of distribution of tdTomato expression indicated notable differences in the access of the two AAV serotypes to primary afferent neurons via peripheral delivery in the colon and to spinal projections neurons via intracranial delivery within the parabrachial complex. Additionally, our results highlight the superior sensitivity of detection of neuronal transduction based on reporter expression relative to expression of viral products.

## INTRODUCTION

AAV vectors have been used for transgene delivery to different components of the nociceptive system in rodents for nearly two decades. Transduction of primary afferent neurons has been demonstrated using intrathecal [1, 2], intraganglionic [3–5], intraneural [6, 7], or peripheral [8, 9] injections. Intraspinal AAV vector delivery has substantially aided neuroanatomical and functional investigations of dorsal horn circuits (for example, [10–12]). In addition, intraparenchymal AAV injections within various brain nuclei have been employed in mapping of supraspinal circuits involved in pain processing [13–15].

AAV serotypes vary in their transduction efficiency via different routes of administration. A number of serotypes have been characterized for intraganglionic, intraspinal, intrathecal, and to a lesser degree peripheral delivery [7, 16, 17]. Among these, AAV9 is distinguished by its efficient transduction of DRG neurons following intrathecal delivery [1] and its capacity for neuronal transduction following localized peripheral delivery [18]. AAV serotypes also differ in their ability to enter neurons via their axon terminals and undergo retrograde transport to the cell body, a property that is particularly useful for targeting of projection neurons in the central nervous system [19]. AAV2retro is a recombinant virus that was specifically engineered for retrograde neuronal transduction [20], and its use has contributed extensively to mapping of neural circuits (for example, see [21–24]).

The main objective of this study was to explore AAV2retro-mediated gene transfer within the nociceptive system and compare it to that mediated by AAV9 [19]. In addition to the widely used intrathecal and intraspinal routes of delivery, we characterized transduction of spinal projection neurons via viral delivery of AAV9 and AAV2retro to the parabrachial nuclear complex. We further employed intracolonic AAV9 and AAV2retro injections to evaluate the potential for selective gene transfer to DRG neurons based on site-specific peripheral viral vector delivery. Neuronal transduction by AAV9 or AAV2retro, driving the expression of a fusion Cre/GFP protein, was evaluated based on Cre-dependent expression of tdTomato in transgenic reporter mice. Although we did not conduct direct quantitative comparisons between the two serotypes due to differences in the viral preparations, our observations indicate differences in the access of AAV9 and AAV2retro to neurons of the nociceptive system that will inform future applications of these vectors.

## METHODS

### Animals

All procedures were performed in accordance with the National Institutes of Health *Guide for the Care and Use of Laboratory Animals* and approved by the University of Minnesota Institutional Animal Care and Use Committee. Experiments were performed in male and female transgenic mice harboring a flox-stop-tdTomato reporter gene (Ai14 mice; jax# 007914) and in wild type C57/Bl6 mice. Mice were 4-10 weeks old at the time of viral injections.

### Viral Vectors

AAV2retro.*h.Syn*.Cre-eGFP, abbreviated as AAV2retro was packaged by the Viral Vector and Cloning Core (VVCC) at the University of Minnesota. AAV9.*h5yn*.Cre-eGFP, abbreviated as AAV9, was packaged by the VVCC or obtained from the University of Pennsylvania Vector Core (see below for titer information). AAV9 and AAV2retro contained the same construct (addgene plasmid #105540, courtesy of Dr. James Wilson), which encodes a Cre/GFP fusion protein under control of the human synapsin promotor. Thus, viral transduction leads to Cre-mediated recombination and tdTomato expression in transduced neurons of Ai14 mice.

### Parabrachial Nucleus Injections

Mice were mounted in a stereotaxic frame (Stoelting, 51600, Wood Dale, IL) under isoflurane anesthesia (2.5-4%) and injected with 500 nL of AAV9 (3×10^13^ GC/mL), AAV2retro (2.5×10^14^GC/mL), or Fast-DiI (2.5 mg/ml in dimethylformamide; ThermoFisher, Waltham, MA). A pulled borosilicate glass pipette with a tip diameter of ~20 μm was attached to a picospritzer (Parker Hannifin, Hollis, NH) and a microsyringe (Hamilton, Reno, NV) by way of a three-way stopcock and the dead space was filled with mineral oil. The pipette was attached to a stereotaxic manipulator (Stoelting) and the tip was advanced 3 mm ventral from the level of the meninges through a small burr hole in the skull positioned 1.5 mm left of midline at a location 0.5 mm caudal from true lambda. The injectate was delivered under pressure, the pipette was slowly removed from the brain after a 5min rest period, and the skin was closed with VetBond (3M, St,Paul, MN). Animals were allowed to recover on a heated pad, and meloxicam (2 mg/kg, s.c.) was given for post-operative analgesia. Tissues were collected for analysis 6 (AAV) or 4 (DiI) weeks after injections.

### Intraspinal injections

Mice were anesthetized with a mixture of ketamine/xylazine/acepromazine (75mg/kg ketamine, 5mg/kg xylazine, 1 mg/kg acepromazine, i.p.). The back skin of anesthetized animals was incised and the soft tissue overlying the L3-L4 spinal cord segments was blunt dissected to the level of the vertebral laminae. The ligament between the T12 and T13 laminae and the underlying dura mater were parted and the right dorsal horn was injected with 1 μL of AAV9 (3×10^13^ GC/mL, n = 3; 8.5×10^13^ GC/mL, n = 3), or AAV2retro (2.5×10^14^GC/mL; n = 4), using the same pressurized pipette setup as for PBN injection. The muscles and subcutaneous layers were then reapproximated and closed with surgical suture (4-0 monocryl with PC-5 needle), and the skin was closed with surgical staples. Animals were allowed to recover on a heated pad and meloxicam (2 mg/kg, s.c.) was given for post-operative analgesia. Tissues were collected for analysis 2 weeks after injections.

### Intrathecal injections

AAV9 (8.5×10^13^ GC/mL) or AAV2retro (1×10^13^GC/mL) were delivered intrathecally (i.t.) by direct lumbar puncture in conscious adult mice as described [13,21]. The mice were gently gripped by the iliac crest, and a 30-gauge, 0.5-inch needle connected to a 50 μL Luer-hub Hamilton syringe was used to deliver 10 μL of injectate into the subarachnoid space over 1-3s at the level of the cauda equina between the L5/L6 vertebrae. The duration of the procedure was approximately 15-30 s per mouse. All injections were confirmed by observation of a tail flick upon entry into the intrathecal space. Tissues were collected for analysis 3 weeks after injections.

### Intracolonic injections

Intracolonic injections were performed in two cohorts of Ai14 mice: cohort 1 received AAV9 (2.7×10^13^ GC/mL; n = 3) or AAV2retro (2.5×10^14^GC/mL; n = 3); cohort 2 received AAV9 (8.5×10^13^ GC/mL; n = 5) or AAV2retro (1×10^13^GC/mL; n = 6). Based on tdTomato labeling in the DRG, the injection success rate for cohort 1 was 100% and for cohort 2 ~50%. In addition, three C57/Bl6 mice were injected with the AAV9 vector. Animals were anesthetized using isoflurane (2.5-4%). A vertical incision (~ 3 cm) was made in the lower abdomen of adult mice, and the descending colon was exposed as described [25]. Mice received a single 4-μl injection of viral vector. Injections were made into the wall of the descending colon horizontal to the longitudinal axis of the colon, using a 10-μl Hamilton syringe attached to a 30G needle. The needle was left in place for 1 minute to prevent reflux. Following injection, the abdominal wall was sutured (Ethicon Cat# Z304H), and the overlying skin was closed with Vetbond. The mice were allowed to recover on a heated pad and meloxicam (2 mg/kg, s.c.) was given for post-operative analgesia. Tissues were collected for analysis 3-4 weeks after injections.

### Immunohistochemistry

Mice were deeply anaesthetized and perfused via the heart with calcium-free Tyrode’s solution (in mM: NaCl 116, KCl 5.4, MgCl_2_·6H_2_0 1.6, MgSO_4_·7H_2_O 0.4, NaH_2_PO_4_ 1.4, glucose 5.6, and NaHCO_3_ 26) followed by fixative (4% paraformaldehyde and 0.2% picric acid in 0.1M phosphate buffer pH 6.9 or 4% paraformaldehyde in phosphate buffered saline). Tissues were dissected and incubated in 10% sucrose (in PBS) solution overnight. Tissues were then cryostat-sectioned at 14-30μm, and the sections were mounted onto gel-coated slides and stored at −20° C until use. Sections were thawed and incubated in diluent (PBS with 0.3% Triton-X100; 1% BSA, 1% normal donkey serum) for 1h at room temperature followed by incubation in primary antibodies (rabbit anti Ds-red, Takara, cat# 632496, 1:1000; or chicken anti-GFP; Abcam, cat# ab113970, 1:1000) overnight at 4° C. Sections were rinsed in PBS (3×10 min each), incubated in species-appropriate secondary antibodies (Cy3-conjugated donkey anti-rabbit, 1:300, cat# 711-165-152, and Alexa488-conjugated donkey anti-chicken, 1:100, cat# 703-545-155, Jackson ImmunoResearch, West Grove, CA) for 1h at room temperature, rinsed again using PBS, and coverslipped using glycerol and PBS containing p-phenylenediamine (Sigma). The specificity of the antisera was confirmed by lack of labeling in the absence of viral injections. Following incubation in primary and secondary antibodies, some DRG sections were stained in NeuroTrace (1:1000, Invitrogen). For visualization of Fast-DiI, tissue sections were rehydrated in diluent for 30 min and coverslipped.

### Imaging and quantification

Images were acquired in an Olympus BX2 microscope equipped with a Fluoview 1000 scan head, software version 4.1.5.5; objectives - UPLAPO 4x/0.16 NA (Fig. 1C1), UPLSAPO 20x/0.85 NA (Fig. 1 A2, B2, C2; Fig. 4 D-F), UPLAPO 10x/0.4 NA (Fig. 2, Fig. 3, Fig. 4 A2, B2); confocal aperture was set to software-determined auto settings. Single or sequential multi-fluorescent images were collected using 405 nm (NeuroTrace), 488 nm (Alexa488), and 543 nm (Cy3 or DiI) laser excitation and collecting emission between 425-475 nm, 505-525 nm, and 550-650 nm, respectively. Images we also collected with a Nikon FN1 upright stand equipped with an A1R HD MP laser scanning head and a motorized Prior stage and piezo Z drive (for sample positioning and focus) and controlled with NIS Elements 5.1 software; objectives - Nikon Plan Fluor DIC 10x/0.3NA (Fig. 1A1, 1B1, 1F; pinhole 12.77 μm, confocal mode, excitation 561 nm, emission 575-625 nm; tiled 4×4 acquisition for A1 and B1) and Nikon Plan Apo LWD 25x water-immersion/NA 1.1 (Fig. 4A1, 4B1; multiphoton mode, Mai Tai DeepSee 920 nm excitation, 570-640 nm emission, GaAsP NDD). For optimal visualization, the representative images used in the figures were adjusted for brightness, contrast, and color using Adobe Photoshop software. Non-linear adjustment was applied simultaneously to images from the same experiment.

**Figure 1.**
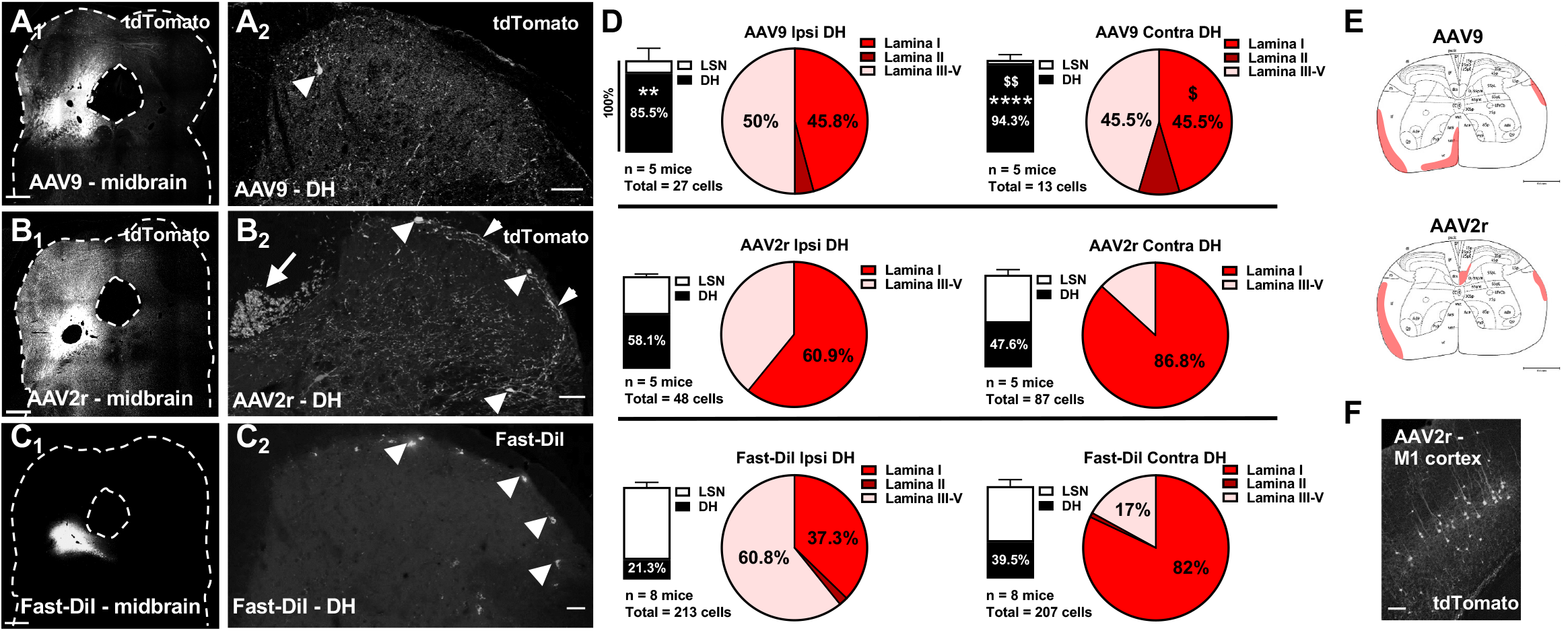
Targeting of spinoparabrachial neurons by AAV9, AAV2retro, and DiI. **A-C,** Representative images of the parabrachial complex (A1, B1, C1; scale bars = 500 μm) and dorsal spinal cord (A2, B2, C2; scale bars = 50 μm) following parabrachial complex injections of AAV9 (A), AAV2retro (AAV2r, B), or Fast-DiI (C). The images in B2 and C2 represent a maximum intensity projection of 3D stacks spanning 12 and 16 μm, respectively, and collected with a 4-μm z-step. Arrowheads in A2, B2 and C2 highlight labeled spinoparabrachial neurons in the superficial dorsal horn. Additionally, in B2, the short arrow highlights a plexus of labeled processes in lamina I, and the long arrow indicates labeling of the dorsal corticospinal tract. **D,** Stacked Bars represent the relative distribution of labeled spinoparabrachial neurons between the dorsal horn (DH) and the lateral spinal nucleus (LSN) in the ipsilateral (left) and contralateral (right) spinal cord following injection with AAV9 (top), AAV2retro (middle), or Fast-DiI (bottom) (** p<0.01, **** p<0.0001, compared to Fast-DiI; $$ p<0.01, compared to AAV2r). The total number of labeled neurons analyzed and the number of animals (n) is noted under the stacked bars. Pie charts in D show the percentage of labeled neurons present in lamina I-V of DH. Proportions of labeled neurons in spinal laminae were compared using Two-way ANOVA and Bonferroni posttest ($, p<0.05 compared to AAV2r). **E,** The average labeling of spinal tracts at the level of L3 is shaded for AAV9 (E1) and AAV2r (E2). **F,** Retrograde labeling of cortical neurons in primary motor cortex (M1) following parabrachial injection of AAV2retro; scale bar = 150μm.

**Figure 2.**
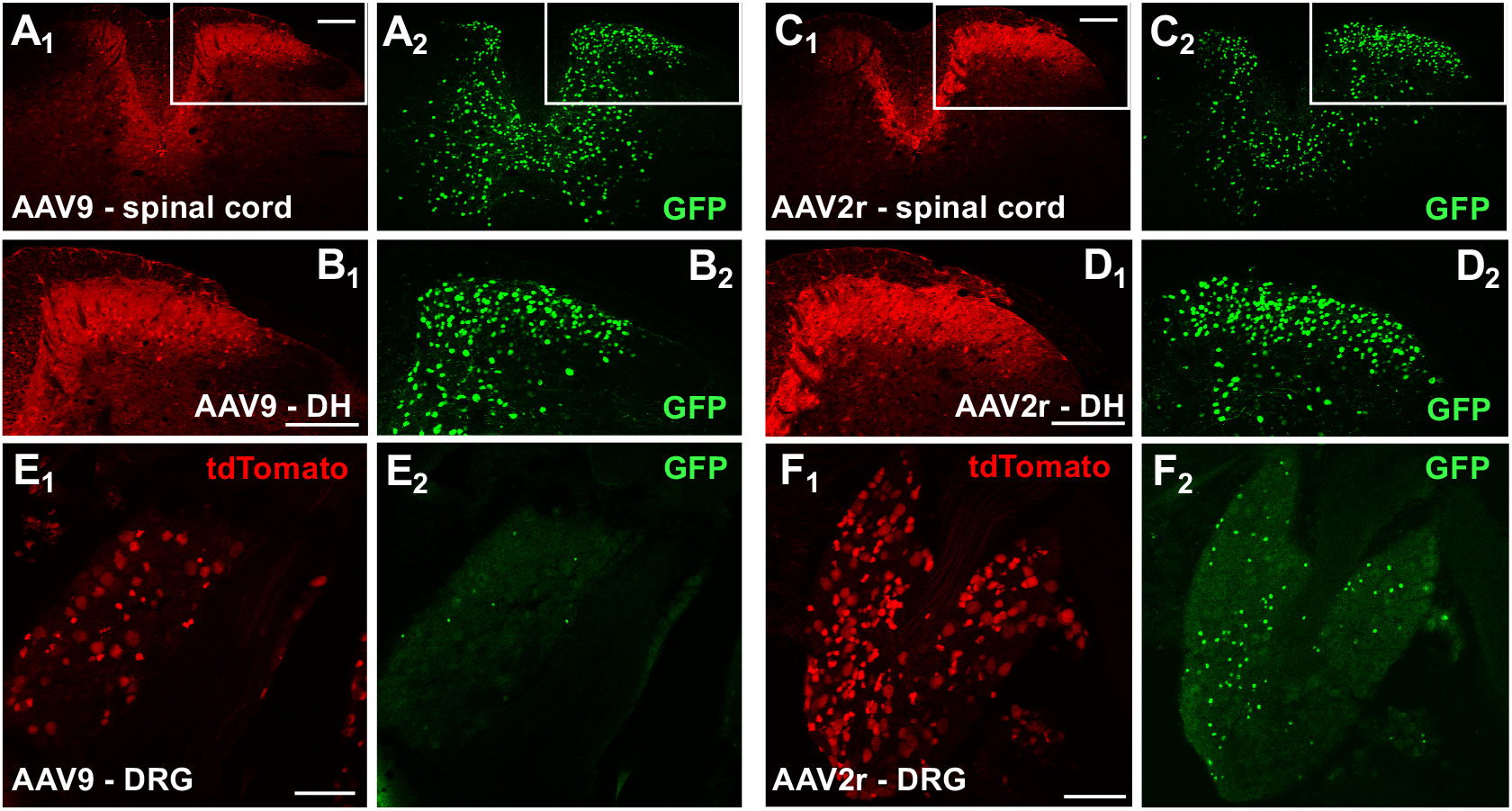
Intraspinal delivery of AAV9 and AAV2retro viral vectors leads to effective transduction of both spinal and primary afferent neurons. **A-D,** tdTomato (red, subscript 1) and GFP (green, subscript 2) immunofluorescence in Ai14 mice injected unilaterally in the L3/L4 region of spinal cord with AAV9 (A and B), or AAV2retro (C and D). **E and F**, tdTomato (red, subscript 1) and GFP (green, subscript 2) immunofluorescence in neurons of the ipsilateral L4 DRG following transduction with AAV9 (E) or AAV2retro (F); not all tdTomato-labeled cells were immunoreactive for nuclear GFP. Scale bars = 150μm.

**Figure 3.**
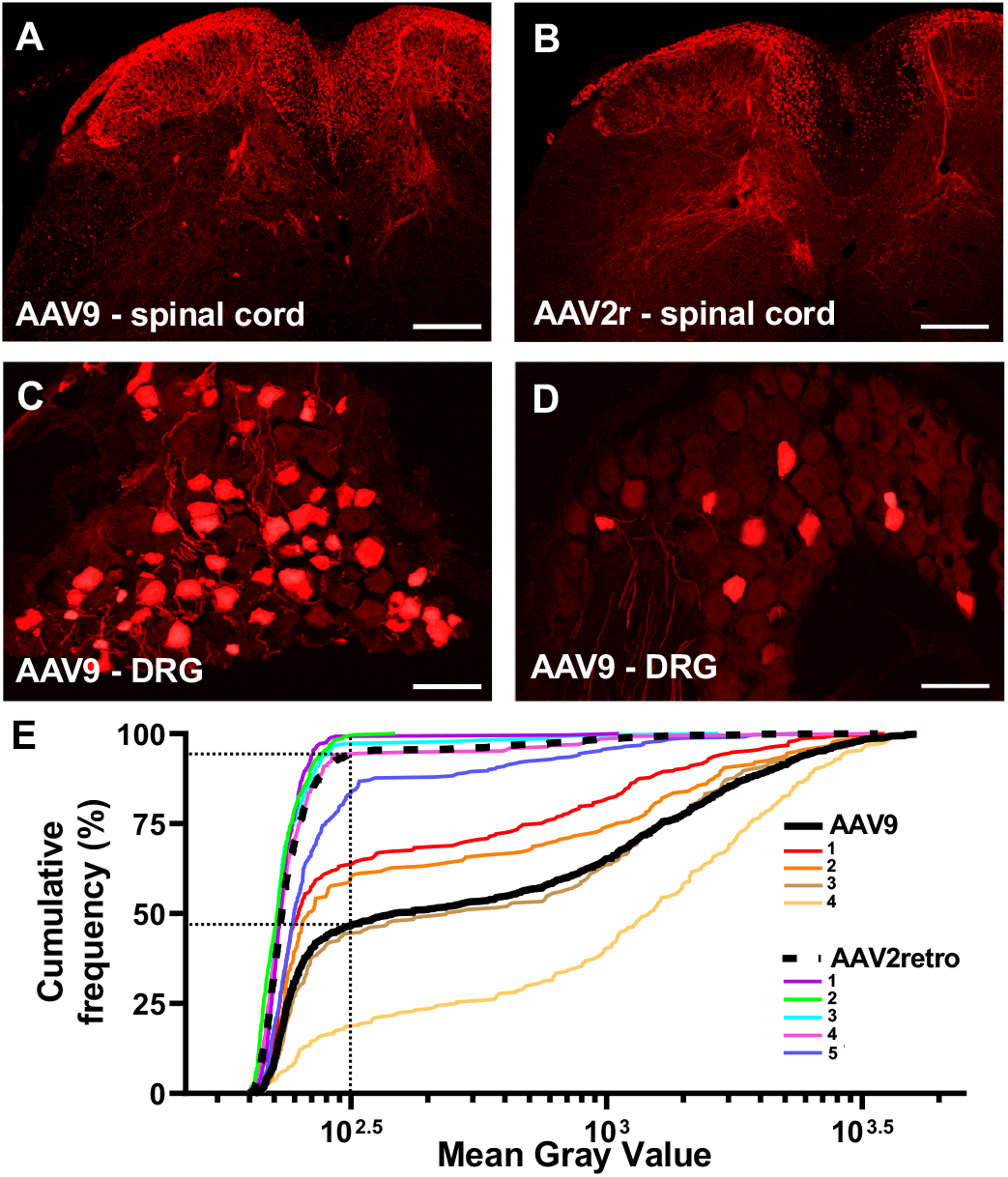
Transduction of spinal cord and DRG neurons following intrathecal delivery of AAV9 and AAV2retro. **A and B,** tdTomato immunofluorescence in the lumbar spinal cord of mice injected with AAV9 (A) or AAV2retro (B); scale bars = 200μm. **C and D,** Transduced neurons in L4 DRG following injection of AAV9 (C) or AAV2retro (D); scale bars = 100μm. **E**, The mean gray value of tdTomato immunofluorescence in individual DRG neurons sampled from L4 DRG of mice injected with AAV9 or AAV2retro is shown as a relative cumulative frequency distribution for individual animals (color lines). The group relative cumulative distributions for individual neurons samoled from all AAV9- and AAV2retro-treated L4 DRG are indicated by a solid black line (AAV9, 997 cells) and a dashed black line (AAV2retro, 1440 cells) and show that more neurons within AAV2retro-treated DRG (~90%) have low mean gray values than neurons within AAV9-treated DRG (<50%) (see vertical and horizontal dotted lines; Kolmogorov-Smirnov test, p<0.001).

**Figure 4:**
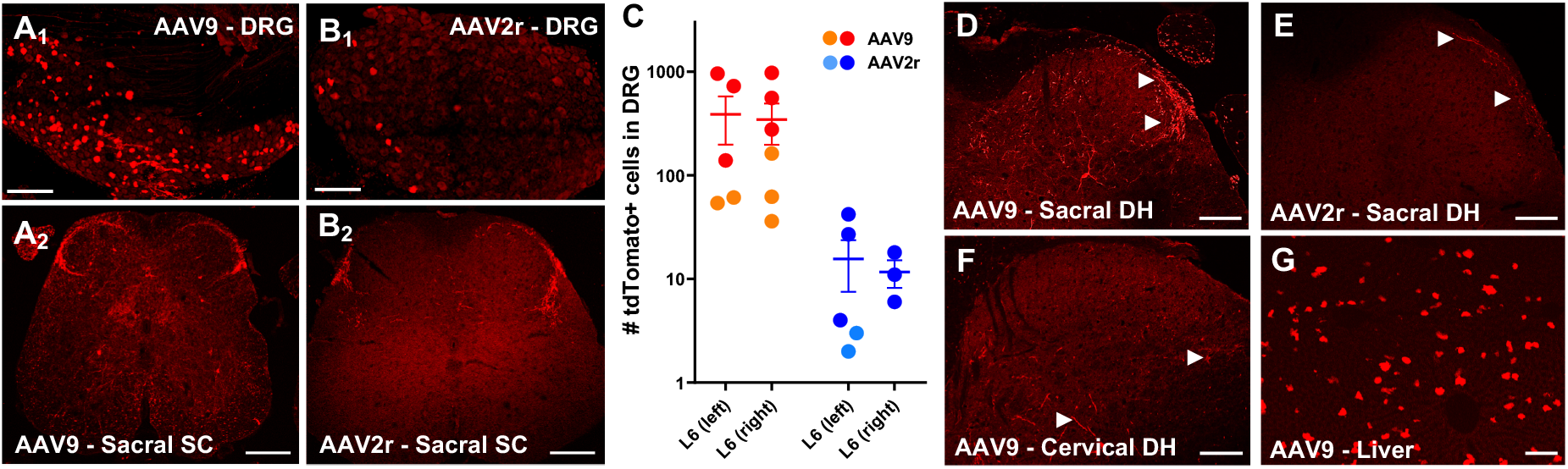
AAV9 and AAV2retro-mediated transduction following intracolonic injections. **A and B**, tdTomato labeling in L6 DRG and sacral spinal cord was more abundant in AAV9-injected compared to AAV2retro injected animals from cohort 1. The images in A1 and B1 represent a maximum intensity projection of four stitched 3D stacks spanning the entire ganglion (collected with a 3 μm z-step). Scale bars: A1 and B1, 150 μm; A2 and B2, 100 μm. C, Quantitative analysis of tdTomato expressing L6 DRG neurons from cohort 1 (circles) and cohort 2 (squares). D-E, Sacral spinal cords from cohort 2 showed labelling of afferent fibers and a few neurons in the spinal cord, consistent with the previously observed pattern (D, E). Scale bar = 100μm. F-G, The presence of tdTomato labelled fibers and cells in cervical spinal cord as well as cells of the liver provides evidence for systemic distribution of AAV9 (F, G). Scale bars: F, 100μm; G, 150μm.

Spinal neurons labeled by parabrachial injections in the rostral lumbar enlargement (L1-L3) were counted manually and mapped by a blinded observer under wide field epifluorescence in 18 30-μm immunostained tissue sections separated by 120 μm. Brains were sectioned coronally at 30μm and sections were immunostained and assessed for tdTomato immunolabeling by a blinded observer at intervals of 600 μm.

Different approaches for quantitative analysis of transduction in DRG neurons were applied for intrathecal and intracolonic AAV delivery to allow comparisons with previous literature. For intrathecal AAV delivery, DRG sections were immunolabeled and counterstained with NeuroTrace (1:1000, Invitrogen). Confocal images were collected with the Olympus Fluoview system as described above (UPLAPO 10x/0.4 NA), using uniform imaging parameters that avoided saturation. Image analysis was performed by trained observers blinded to the experimental groups using Fiji. NeuroTrace-labeled neuronal profiles with nuclei were outlined in 5 evenly spaced sections per DRG, and the mean grey value of tdTomato immunoreactivity in each profile was measured. To estimate the proportion of labeled neurons, a threshold mean gray value was selected for the entire data set based on the rate of change of the slope of the cumulative mean gray value frequency distribution. For quantitative analysis of transduction in DRG neurons after intracolonic AAV delivery, native tdTomato and GFP fluorescence was imaged in whole-mounted L6 DRG (without clearing) on a two-photon Nikon A1R system as described above for Fig. 4A1 and 4B1. tdTomato or GFP-expressing neurons in the z-stacks were manually counted by an observer blinded to the treatments.

### Statistics

The number of animals in each condition was used as the value of n. The distribution of spinoparabrachial neurons within the dorsal spinal cord was compared by two-way ANOVA with Bonferroni corrected post testing for pairwise comparisons, α = 0.05 (AAV treatment x spinal laminae, data shown in Fig. 1 pie charts). Comparisons between ipsilateral and contralateral distributions were made using Fisher’s exact test (data shown in Fig. 1 stacked columns and pie charts). For intrathecal injections, the relative cumulative distributions of the mean gray values of labeled DRG neurons were compared using the Kolmogorov-Smirnov test (Fig. 3E). Statistical analysis and graphical representation of data was performed using Prism 8 software.

## RESULTS

### Differential targeting of spinoparabrachial neurons by AAV9 and AAV2retro

To selectively transduce spinoparabrachial neurons, Ai14 reporter mice were injected in the parabrachial complex with AAV9 (3×10^13^ GC/mL) or AAV2retro (2.5×10^14^GC/mL) vectors (n = 5 per treatment) expressing GFP-tagged Cre recombinase. To compare viral transduction rates with a retrograde labeling method employed in functional assays, a separate cohort of C57/Bl6 mice (n = 8) was injected with the lipophilic fluorescent dye Fast-DiI. Figure 1 shows examples of parabrachial injection sites and labeled spinal projection neurons in histological sections derived from Ai14 mice injected with AAV9 (Fig. 1A), AAV2retro (Fig. 1B), or Fast-DiI (Fig, 1C). Expression of the tdTomato reporter protein and the viral Cre-GFP was visualized by immunolabeling. Notably, GFP labeling was rarely observed in the tdTomato-positive transduced neurons, suggesting that the expression levels Cre-GFP were below the detection limit of visualization, although sufficient for Cre-dependent recombination. Ai14 mice without viral injection showed no tdTomato fluorescence in either the brain or the spinal cord (not shown).

The number and location of spinoparabrachial neurons labeled by tdTomato expression or Fast-DiI was quantified in the dorsal horn (DH) and lateral spinal nucleus (LSN) from 30μm transverse sections of the rostral lumbar enlargement (L1-L3). The relative proportions of labeled neurons in the LSN and DH following AAV9 and AAV2retro parabrachial injections was notably different (Fig. 1D, stacked bars). Following AAV9 treatment, over 80% of the retrogradely labeled projection neurons were located in DH. In contrast, AAV2retro labeled projection neurons in the DH and LSN equally. These distinct patterns of labeling were observed in spinal cord both ipsilateral and contralateral to the site of parabrachial injection. The distribution of DiI-labeled neurons largely paralleled that of AAV2retro-targeted neurons, although there was a trend toward higher proportions of DiI-labeled neurons within the LSN, particularly on the ipsilateral side.

The lamina I transduction rates expressed as neurons per 1 mm of rostro-caudal spinal cord length contra- and ipsilateral to the injection site respectively were as follows: 2.5 ± 1.3 and 4.6 ± 2.5 for AAV9, compared to 23.5 ± 8.1 and 7.4 ± 3.3 for AAV2retro. The corresponding labeling rate for Fast-DiI was 42.7 ± 16.8 in the contralateral lamina I, and 9.9 ± 3.9 on the ipsilateral side. Although differences in vector titer precluded direct comparison of transduction efficiency between the AAV9 and AAV2retro vectors, we compared the distribution of labeled DH neurons between the treatment groups (Fig. 1D, pie charts). The proportion of AAV9 labeled projection neurons found in lamina I of the contralateral DH was significantly lower than that observed with AAV2retro, which showed a pattern similar to Fast-DiI. Furthermore, the proportions of labeled neurons present in lamina I versus deep laminae III-V was significantly different between ipsilateral and contralateral sides of animals injected with AAV2retro (Fisher’s exact test; p<0.05) or Fast-DiI (Fisher’s exact test; p<0.0001). This pattern was not observed with AAV9, which showed no significant difference in the relative distribution of labeled neurons between the ipsilateral and contralateral side (Fisher’s exact test; p>0.05).

AAV2retro was readily transported retrogradely and transduced neurons in numerous distant areas of the brain including cingulate cortex, orbital cortex, primary motor cortex and primary somatosensory cortex. Consistent with the transduction of neurons in primary motor cortex (Fig. 1F), we observed prominent labeling of the dorsal corticospinal tract (Fig. 1 B2, long arrow). In contrast, AAV9-mediated transduction was largely limited to the brainstem and was never seen in cortical regions. The labeling of tracts within the white matter of the spinal cord is summarized in Fig. 1E.

### AAV9 and AAV2retro vectors effectively transduce spinal dorsal horn and dorsal root ganglion neurons when delivered intraspinally

To assess the utility of AAV9 and AAV2retro in targeting the somatosensory system at the level of the spinal cord, we performed intraspinal injections of the vectors into the lumbar enlargement of Ai14 mice (AAV9, n = 3; AAV2retro, n = 4) and observed viral transduction of neurons in the spinal cord and dorsal root ganglia (DRG). Figure 2 shows representative images of immunolabeling of the tdTomato reporter and the virally expressed Cre-GFP in spinal cord from mice injected with AAV9 (Fig2. A & B) or AAV2retro (Fig2. C & D). The highest levels of labeling for both tdTomato and GFP were seen in the ipsilateral dorsal horn. In this region, tdTomato labeling was noted both in neuronal somata and in their processes, whereas GFP labeling was localized in neuronal nuclei consistent with the nuclear localization of Cre. tdTomato-expressing neuronal somata were also evident in the L4 DRG ipsilateral to the intraspinal injections of AAV9 (Fig. 2E) and AAV2retro (Fig. 2F). The level of GFP expression in the DRG as compared to the dorsal horn was consistently lower. Furthermore, the virally expressed GFP was not detected in a large portion of tdTomato-positive DRG neurons. These data show that both AAV9 and AAV2retro effectively transduce somatosensory neurons of the dorsal horn and DRG when injected intraspinally. In addition, they highlight the higher sensitivity in detecting AAV transduction based on reporter expression (tdTomato) compared to viral protein expression (Cre/GFP).

### AAV9 and AAV2retro vectors transduce DRG neurons following intrathecal delivery

Intrathecal injection offers the least invasive route of viral vector administration to the CNS. Intrathecal delivery of 10 μl of AAV9 (8.5×10^13^ GC/mL, n = 5) or AAV2retro (1×10^13^ GC/mL, n = 6) resulted in prominent tdTomato labeling in spinal dorsal horn (Fig. 3A and 3B). Compared to intraspinal viral vector delivery (Fig. 2), relatively few spinal neurons were transduced, and the vast majority of tdTomato immunofluorescence present in the spinal cord was localized to the central processes of primary afferent neurons (identified morphologically). This was confirmed by a relative dearth of viral GFP immunofluorescence in spinal cord (not shown), and expression of tdTomato by transduced DRG neurons (Fig 3C-E). Estimates of proportions of viral labeled neurons indicated that approximately 54% (range 36 – 82%) of L4 DRG neurons were transduced by intrathecal delivery of AAV9, while AAV2retro delivery transduced approximately 6% (range 0.7 −20%). The difference in the number of AAV9- and AAV2retro-transduced DRG neurons is also illustrated by the relative cumulative frequency distribution of the mean gray values of tdTomato labeling in DRG neurons (Fig. 3E).

### AAV9- and AAV2retro transduce DRG neurons differentially via their peripheral processes

To determine whether AAV9 and AAV2retro transduce primary afferent neurons via their peripheral processes, we examined transduction in DRG following delivery of the vectors into the wall of the descending colon. In the mouse, the sensory innervation of the descending colon is provided predominantly by L5, L6, and S1 DRG [26]. Transduction was evaluated based on detection of native tdTomato and GFP fluorescence within DRG neurons in whole-mount preparations of L6 DRG from Ai14 mice injected with AAV9 or AAV2retro vectors driving expression of Cre-GFP. A single intracolonic injection of AAV9 (2.7×10^13^ GC/mL) yielded a higher number of tdTomato-expressing DRG neurons than an injection of AAV2retro (2.5×10^14^GC/mL), indicating that AAV9 transduces DRG neurons more effectively than AAV2retro via this route of delivery (Fig. 4 A1, B1 and C). Strikingly, GFP fluorescence was not observed in AAV9- or AAV2retro DRG (not shown). To test whether the lack of native GFP fluorescence was entirely attributable to a detection limit in the whole-mount preparations, we examined DRG transduction with the same AAV9 vector in two wild-type C57/Bl6 mice. Counting of labeled cells in two L6 DRG from these mice yielded 132 and 55 GFP-positive neurons, respectively. These results suggest that viral expression of Cre/GFP in DRG neurons may be suppressed following Cre-dependent recombination in Ai14 mice. We are not aware of previous reports of similar observations, and the potential underlining mechanisms are unclear.

At lower lumbar and sacral spinal cord levels, both AAV9 and AAV2retro intracolonic treatments in Ai14 mice resulted in tdTomato labeling of nerve fibers in superficial dorsal horn and to a lesser extent in the intermediate gray matter (Fig. 4 A2, B2, D, and E). Surprisingly, we also observed scattered labeling of neurons within the spinal cords of AAV9-treated mice (Fig. 4 A2 and D). While some of these neurons could be preganglionic parasympathetic neurons projecting to the colon, it is also possible that the labeling of spinal neurons is due to trans-synaptic transfer of AAV9 [27] or to access of AAV9 to the systemic circulation following intracolonic delivery. To address the latter possibility, we treated a second cohort of mice with a different set of AAV9 (8.5×10^13^ GC/mL) or AAV2retro (1×10^13^GC/mL) vectors (same as used for intrathecal injections) and extended the histological analysis to cervical spinal cord (Fig 4F) and liver (Fig. 4G). The superior level of transduction achieved by AAV9 was consistent between the two cohorts (Fig. 4C). In lower lumbar and sacral spinal cord, we observed labeling for both vectors in a pattern similar to the first cohort. In contrast, in cervical spinal cord, tdTomato labeling was observed in AAV9- (Fig. 4F) but not AAV2retro-treated (not shown) mice. Unlike lower lumbar and sacral levels (Fig 4A2), the pattern of AAV9-mediated tdTomato expression in cervical spinal cord consisted of scattered fibers and few neurons without prominent labeling of fibers in superficial dorsal horn (Fig 4F). Transduction was also noted in the liver of AAV9- (Fig. 4G) but not AAV2retro-treated (not shown) mice following intracolonic delivery. The presence of AAV9-mediated transduction in cervical spinal cord and liver suggests that AAV9 gains access to the systemic circulation following intracolonic delivery.

## DISCUSSION

In the present study, we characterized the ability of AAV2retro and AAV9 to transduce components of the nociceptive system via different routes of delivery. Our results indicate notable differences in the access of the two AAV serotypes to primary afferent neurons via peripheral delivery and to spinal projection neurons via intracranial delivery within the parabrachial complex. These differences likely result from mechanisms that govern the diverse cellular tropism of AAV serotypes and remain poorly understood [28]. AAV9 and AAV2 (the parent serotype of AAV2retro) bind to different primary and secondary receptors on the cell surface [28]. AAV9 is distinguished from other serotypes by its more efficient systemic distribution to various tissues and organs and its ability to cross the blood-brain barrier [1, 29]. On the other hand, AAV2retro harbors engineered capsid mutations that, through unknown mechanisms, confer enhanced transduction efficiency via retrograde transport [20].

We evaluated neuronal transduction based on Cre-dependent expression of tdTomato, controlled by the CAG promoter, following delivery of AAV9 or AAV2retro carrying identical constructs that drive the expression of Cre/GFP, under the control of the hSyn promoter. tdTomato and Cre/GFP labeling overlapped extensively in spinal neurons transduced via intraspinal delivery. However, we observed multiple instances of spinal projection neurons and DRG neurons that were tdTomato-positive but lacked Cre/GFP labeling, indicating that Cre/GFP expression levels were sufficient for Cre-recombination but below the detection limit for immunohistochemical detection. The discrepancy in tdTomato and Cre/GFP labeling in these neurons may reflect entry of fewer viral particles and the fact the CAG promoter is substantially stronger than the hSyn promoter. These observations highlight the superior sensitivity of detection of neuronal transduction based on reporter expression (i.e. tdTomato) relative to expression of viral products (i.e. Cre/GFP). Limitations in the histological detection of viral transgene expression may affect data interpretation and the assessment of off-target effects in functional studies.

### Transduction of spinal projection neurons

The spinal dorsal horn functions as the initial site of integration between sensory information emanating from peripheral tissues and descending modulation from supraspinal levels. Spinal projection neurons constitute only 1% of neurons in the dorsal horn and serve as the output of this integrative circuit. Projection neurons located in superficial lamina I tend to display a high threshold “nociceptive specific” phenotype. In rodents, the vast majority of these nociceptive-specific projection neurons project to the parabrachial nucleus in the midbrain [30]. We targeted these cells for genetic manipulation with viral vectors to aid future investigation into their function and connectivity.

We used the lipophilic dye Fast-DiI as a “universal” retrograde tracer of spinoparabrachial neurons. Although Fast-DiI has been employed for the identification of spinoparabrachial neurons in functional studies [31–34], to our knowledge, the spinal distribution of labeled neurons has not been evaluated histologically in the mouse. Previous neuroanatomical analysis of spinoparabrachial neurons in the mouse was conducted using the retrograde tracer cholera toxin B (CTB) [30]. This study reported an approximately 3-fold higher number of retrogradely labeled neurons at the L4 spinal level than the number of Fast-DiI neurons we observed within the rostral (L1-L3) lumbar spinal cord. This difference in the rate of labeling of spinoparabrachial neurons may be due to the use of different retrograde tracers and the distinct anatomical locations of the analyses.

Comparison of the distribution of spinoparabrachial neurons labeled by Fast-DiI, AAV2retro and AAV9 suggests marked differences. Most notably, Fast-DiI was more likely to label projection neurons in the LSN than the AAV vectors. The observation of LSN spinoparabrachial neurons here is consistent with recent identification of a sustained pain coping pathway involving LSN neurons that project to the parabrachial nucleus in the mouse [35]. Our results suggest that AAV9 and AAV2retro may exhibit lower tropism for LSN compared to dorsal horn spinoparabrachial neurons.

Treatment with both AAV2retro and Fast-DiI yielded more labeled neurons in lamina I of the contralateral than the ipsilateral dorsal horn. This is consistent with reports showing that the majority of lamina I spinoparabrachial neurons project to the contralateral midbrain. However, this pattern of labeling was not seen following AAV9 treatment. The different pattern of transduction of spinoparabrachial neurons by AAV9 and AAV2retro may reflect differential tropism of the two serotypes for different subsets of projection neurons under our experimental conditions. Interestingly, a recent study reported that lamina I spinoparabrachial neurons were efficiently transduced by intraparabrachial complex injection of AAV9, driving expression of the calcium indicator GCamP6s under the control of the strong CAG promoter [36]. The difference in the strength of the CAG and hSyn promoters likely accounts for the difference between these observations and our results, consistent with a previous report [16].

### Transduction of dorsal horn and DRG neurons following intrathecal and intraspinal AAV delivery

Transduction of dorsal horn neurons by several AAV serotypes delivered intraspinally has been previously described [16]. Our observations of extensive transduction of dorsal horn neurons by AAV9 are consistent with these findings. We also observed abundant spinal transgene expression in AAV2retro-treated mice, which was qualitatively similar to AAV9-driven expression. This is also consistent with the findings of Haenraets at al. (2017), where AAV serotypes 1, 6, 7, 8, 9, and 10 displayed similar transduction efficiencies in dorsal horn. In contrast to intraspinal delivery, we observed limited transduction of dorsal horn neurons via intrathecal delivery; this observation is consistent with previous reports for AAV9 and other serotypes carrying single-stranded viral constructs [1, 2, 37].

We found that both AAV9 and AAV2retro transduced DRG neurons following intraspinal and intrathecal delivery, although differences in viral titers precluded quantitative comparisons of transduction efficiency. Considering several methodological differences, the transduction efficiency we observed for intraspinally delivered AAV9 (~21% of L4 DRG neurons) is consistent with the findings of Haenraets et al. (2017). Although we found that the minimally invasive intrathecal injection of AAV9 transduced a higher proportion of L4 DRG neurons (54%, range 36-82%) than intraspinal, it should be noted that the dose administered intrathecally was ten-fold higher. In addition, while DRG transduction following intraspinal viral delivery is likely restricted to several segments, transduction via intrathecal delivery is more broadly distributed [1, 38]. Thus, the two routes of administration offer a number of advantages and disadvantages that can be tailored to specific experimental requirements.

### Transduction of DRG following intracolonic delivery of AAV

Our observations indicate that AAV9 transduces DRG neurons more efficiently than AAV2retro following intracolonic injection and that AAV9 also appears to gain access to the systemic circulation. The latter conclusion is based on the presence of tdTomato labeling in liver and cervical spinal cord. However, the distinct pattern of labeling in superficial lumbosacral and cervical dorsal horn of AAV9-injected mice suggests that systemic redistribution of the virus accounts for only a small portion of the transduction observed in L6 DRG. Delivery of lower injection volumes in the periphery may reduce systemic redistribution of AAV9 and improve the selective targeting of neurons innervating the injection site. The extent of transduction due to access of AAV9 to the systemic circulation may also be sex dependent as there is evidence for differential transduction of CNS and liver of male and female mice following intravenous AAV9 delivery [39, 40]. Finally, it remains to be determined whether intracolonic delivery of AAV9 leads to transduction of preganglionic parasympathetic neurons or trans-synaptic anterograde gene transfer to spinal neurons in lower lumbar and sacral spinal cord.

We observed that AAV2retro intracolonic injection yielded a substantially lower number of transduced DRG neurons, regardless of titer differences of the viral preparations. This suggests that the determinants that confer enhanced transduction efficiency of AAV2retro via retrograde transport in the CNS are not present in the peripheral processes of DRG neurons innervating the distal colon. It is unclear if other peripheral DRG targets would display similar characteristics. However, since AAV2retro-mediated transduction remained limited to the injection site, this serotype may be useful for selective organ-specific targeting of DRG neurons in paradigms where a relatively lower rate of transduction is acceptable.

### Limitations

Several factors that may affect the distribution and/or detection of gene transfer, such as different promoters [16] and survival times, were not examined here, Moreover, different preparations of the same vector may vary substantially [41]. Further investigation of these factors is needed to optimize targeting depending on experimental need.

### Conclusions

The results of the present study characterize the utility of AAV9 and AAV2retro for gene transfer to different components of the nociceptive system. The use of a reporter mouse line allowed for high sensitivity in the assessment of transgene expression and the biodistribution of the viral vectors. Our results suggest differential tropism of AAV9 and AAV2retro for spinoparabrachial neurons. In addition, we observed differences in the targeting of primary afferent neurons via peripheral administration of the viral vector. These observations can assist in tailoring the application of AAV9 and AAV2retro in studies of the nociceptive system.

## AUTHOR’S NOTE

Dr. Skorput’s current affiliation is Molecular and System Biology, Geisel School of Medicine at Dartmouth, Lebanon, NH

## ACKNOWLEDGEMENTS

We are grateful for the resources and expertise by the University of Minnesota University Imaging Centers (UIC, SCR_020997) and especially for the guidance provided by Dr. Guillermo Marques in rigorous reporting of image collection, processing and analysis. We also thank Galina Kalyuzhnaya for technical assistance. The work was supported by NIH grants R34 NS111654 and U18 EB021716.

